# Genome-wide identification of *Aedes albopictus* long noncoding RNAs and their association with dengue and zika virus infection

**DOI:** 10.1101/2020.05.13.093880

**Authors:** Azali Azlan, Sattam M. Obeidat, Kumitaa Theva Das, Muhammad Amir Yunus, Ghows Azzam

**Affiliations:** School of Biological Sciences, Universiti Sains Malaysia, 11800 Penang, Malaysia; Infectomics Cluster, Advanced Medical & Dental Institute, Universiti Sains Malaysia, Bertam, 13200 Kepala Batas, Pulau Pinang, Malaysia

**Keywords:** *Aedes albopictus*, long non-coding RNA, dengue, Zika

## Abstract

The Asian tiger mosquito, *Aedes albopictus* (*Ae. albopictus*), is an important vector that transmits arboviruses such as dengue (DENV), Zika (ZIKV) and Chikungunya virus (CHIKV). On the other hand, long noncoding RNAs (lncRNAs) are known to regulate various biological processes. Knowledge on *Ae. albopictus* lncRNAs and their functional role in virus-host interactions are still limited. Here, we identified and characterized the lncRNAs in the genome of an arbovirus vector, *Ae. albopictus*, and evaluated their potential involvement in DENV and ZIKV infection. We used 148 public datasets, and identified a total of 10, 867 novel lncRNA transcripts, of which 5,809, 4,139, and 919 were intergenic, intronic and antisense respectively. The *Ae. albopictus* lncRNAs shared many characteristics with other species such as short length, low GC content, and low sequence conservation. RNA-sequencing of *Ae. albopictus* cells infected with DENV and ZIKV showed that the expression of lncRNAs was altered upon virus infection. Target prediction analysis revealed that *Ae. albopictus* lncRNAs may regulate the expression of genes involved in immunity and other metabolic and cellular processes. To verify the role of lncRNAs in virus infection, we generated mutation in lncRNA loci using CRISPR-Cas9, and discovered that two lncRNA loci mutations, namely XLOC_029733 (novel lncRNA transcript id: lncRNA_27639.2) and LOC115270134 (known lncRNA transcript id: XR_003899061.1) resulted in enhancement of DENV and ZIKV replication. The results presented here provide an important foundation for future studies of lncRNAs and their relationship with virus infection in *Ae. albopictus*.

**Author summary:** *Ae. albopictus* is an important vector of arboviruses such as dengue and Zika. Studies on virus-host interaction at gene expression and molecular level are crucial especially in devising methods to inhibit virus replication in *Aedes* mosquito. Previous reports showed that, besides protein-coding genes, noncoding RNAs such as lncRNAs are also involved in virus-host interaction. In this study, we report a comprehensive catalog of novel lncRNA transcripts in the genome of *Ae. albopictus*. We also show that the expression of lncRNAs was altered upon infection with dengue and Zika. Additionally, depletion of certain lncRNAs resulted in increased replication of dengue and Zika; hence, suggesting potential association of lncRNAs in virus infection. Results of this study provide a new avenue to the investigation of mosquito-virus interactions that may potentially pave way to the development of novel methods in vector control.

## Introduction

The Asian tiger mosquito, *Aedes albopictus* (*Ae. albopictus*) is an important vector of arboviruses such as dengue virus (DENV), Chikungunya virus (CHIKV), and Zika virus (ZIKV). Due to its invasiveness and aggressive spread, *Ae. albopictus* has widespread geographic distribution; hence, poses serious health threat globally. Although *Ae. albopictus* is considered as a less competent vector of DENV than *Aedes aegypti* (*Ae. aegypti*), *Ae. albopictus* was responsible for dengue outbreaks in Hawaii, China, and Europe, primarily because of its fast expansion across the globe [1]. Due to its epidemiological importance, *Ae. albopictus* has become a key focus for many scientific investigations.

Long noncoding RNAs (lncRNAs) can be defined as RNA species of more than 200 nucleotides (nt) that do not encode protein [2]. Similar to protein-coding mRNAs, lncRNAs are the products of Pol II, and they undergo polyadenylation, capping and alternative splicing [3]. Due to their mRNA-like features, lncRNAs are usually represented in RNA-seq datasets. Once considered as junk and noise in RNA-seq data, findings from many studies provide evidence that lncRNAs perform various biological functions. For example, metastasis associated in lung adenocarcinoma transcript 1 (*Malat1*), is important for alternative splicing of mRNA transcripts, and gene regulation [4], [5]. Besides, research in *Drosophila melanogaster (D. melanogaster)* provide evidence on the functional roles of lncRNAs in spermatogenesis [6] and neurogenesis [7].

Meanwhile, studies in *Ae. aegypti* demonstrated that lncRNAs might be involved in early embryonic development and virus infection [8], [9]. A fraction of *Ae. aegypti* lncRNAs was highly expressed in blood-fed ovaries, where the process of oogenesis takes place. The same set of lncRNAs were also found to be enriched in early embryo (0-8 hour), indicating that they were maternally inherited, and performed critical functions during maternal to zygotic transition stage [8]. Besides, RNAi-mediated silencing of *Ae. aegypti* lncRNA (lincRNA_1317) resulted in enhanced DENV replication in *Ae. aegypti* cells, suggesting the potential role of lncRNAs in virus-host interaction [9]. We speculate that, similar to *Ae. aegypti*, lncRNAs in *Ae. albopictus* may participate in various regulatory functions including virus infection.

Here, we report genome-wide identification of lncRNAs in *Ae. albopictus*, and their possible involvement in DENV and ZIKV infection. In this present work, we defined a high-confident set of 10, 867 novel lncRNA transcripts, derived from 9,298 genomic loci. Our newly predicted *Ae. albopictus* lncRNAs shared similar characteristics with other species such as shorter in length, lower GC content, and lower sequence conservation compared to protein-coding transcripts. To investigate the transcriptional response of lncRNAs upon virus infection, we generated RNA-seq libraries of *Ae. albopictus* cells infected with DENV and ZIKV. Similar to previous studies in *Ae. aegypti* [9], the expression level of *Ae. albopictus* lncRNAs was altered during virus infection. Furthermore, CRISPR-Cas9 induced mutation of selected lncRNAs in *Ae. albopictus* cell led to enhancement of DENV and ZIKV replication, suggesting the potential role of *Ae. albopictus* lncRNAs in antiviral response.

Collectively, the data presented here provide valuable resources for future studies of lncRNAs in *Ae. albopictus* and facilitate a better understanding of the potential role of lncRNAs in virus-host interactions.

## Materials and Methods

### Preparation of public RNA-seq datasets

A total of 148 public RNA-seq datasets were downloaded from NCBI Sequence Reads Archive (SRA) with accession number SRP219966 [10], SRP050258 [11], SRP018112 [12], SRP077936 [13], SRP056407 [14], and SRP164784 [15] All 148 RNA-seq libraries used in this study were listed in **S1 Data**. FASTQC was used to check the quality of raw reads. Adapters and low quality reads (<20 Phred score) were clipped using Trimmomatic version 0.38 [16].

### Mapping and transcriptome assembly

Reads were individually mapped onto *Ae. albopictus* genome (Genome version: AalbF2, assembly: GCA_006496715.1, NCBI) using HISAT2 version 2.1.0 [17]. The resulting alignment files were used as input in Stringtie software (version 1.0.1) for transcriptome assembly [18]. Annotation file (GTF format) of *Ae. albopictus* from NCBI (Aalbo_primary.1_genomic.gtf) was used to guide the transcriptome assembly. A minimum of 200 bp was set for the length of the assembled transcripts. Then, the Stringtie output files were merged into a single transcriptome assembly using Stringtie merge. The resulting merged assembly was then compared to the reference annotation using Gffcompare (https://github.com/gpertea/gffcompare). Transcripts with class code “u”, “i”, and “x” were kept for lncRNA prediction.

### Novel lncRNA prediction

TransDecoder [19] was used to predict the presence of long open reading frame (ORF) within transcripts with class code “u”, “i”, and “x”. A minimum of 100 amino acids was set as threshold of ORF length in TransDecoder analysis, and transcripts having long ORF were removed. The remaining transcripts were then subjected to coding potential assessment analysis. CPC2 [20], PLEK [21], and CPAT [22] were used to evaluate coding potential of the transcripts. For CPAT analysis, a coding probability threshold of < 0.3 was set [8], [9]. Transcripts that passed CPAT threshold, and detected as noncoding by CPC2 and PLEK, were kept for subsequent analyses. To avoid false positive prediction, BLASTX algorithm was used to search for sequence similarity of the potential lncRNA transcripts in Swissprot database. Transcripts with high sequence similarity (E-value < 1e-5) were removed. Transcripts without strand information, and those located less than 1 kb from protein-coding genes on the same strand were removed. Finally, to remove lowly expressed transcripts, transcripts that had an expression of more than 1 TPM in at least 5 RNA-seq libraries were kept. The expression was determined by featureCounts [23], and alignment files (BAM format) generated by HISAT2 were used as input.

### Cell culture and virus

C6/36 cells (ATCC: CRL-1660) were cultured in Leibovitz’s L-15 medium (Gibco, 41300039) supplemented with 10% Fetal Bovine Serum (FBS, Gibco, 10270) and 10% Tryptose Phosphate Broth (TPB) solution (Sigma, T9157). The C6/36 cells were grown at 25°C without CO_2_. BHK-21 cells (ATCC: CCL-10) were used for virus titration. BHK-21 cells were cultured in Dulbecco’s modified Eagles Medium (DMEM, Gibco, 11995065) supplemented with 10% FBS (Gibco, 10270) at 37°C and 5% CO_2_. In this study, DENV serotype 1 (DENV-1) of Hawaiian strain, and ZIKV of strain H/PF/2013 were used. Both viruses were propagated in C6/36 cells. Virus titer of DENV and ZIKV was determined using tissue culture infectious dose 50 (TCID50) assay. DENV-1 (Hawaiian strain) was a gift from David Perera, University Malaysia Sarawak, and ZIKV (H/PF/2013 strain) was a gift from Shee Mei Lok, Duke-NUS Medical School, Singapore.

### Tissue Culture Infectious Dose 50 (TCID50) assay

TCID50 assay was conducted to determine infectious virus titre using a method that has been previously described [24]–[26]. For each determination, a 96-well plate culture of confluent BHK-2 was inoculated with a 150 μl mixture of 100 μl medium and 50 μl of serial 10-fold dilutions of clarified cell culture supernatant media. Each row included 6 wells of the same dilution and two negative controls. Plates were incubated at at 37°C and 5% CO_2_ for 5 days. Each well was observed for the presence or absence of cytopathic effect (CPE). The determination of TCID50/ml was performed using the Reed and Muench method [27].

### Virus infection, RNA extraction and sequencing

C6/36 cells were infected with DENV-1 and ZIKV at a multiplicity of infection (MOI) of 0.25. At 3 days post infection (dpi), the culture medium was removed, and total RNA extraction was carried out using miRNeasy Mini Kit 50 (Qiagen, 217004) according to the manufacturer’s protocol. RNA concentration was determined using ND-2000 instrument (NanoDrop Technologies), and RNA integrity was determined using 2100 Bioanalyzer (Agilent). Libraries were constructed using Illumina TruSeq RNA Sample Prep kit (Illumina, San Diego) using one microgram (μg) of total RNA as starting material. The prepared libraries were sequenced on the Illumina HiSeq 2000 platform to generate 150 bp paired-end reads. Raw reads were deposited in NCBI with accession SRP221722.

### Identification of differentially expressed lncRNAs upon infection

RNA-seq libraries generated in this study were mapped to the *Ae. albopictus* genome (AalbF2, assembly: GCA_006496715.1, NCBI) using HISAT2 [17]. The expression was quantified using featureCounts [23]. GTF file annotated with the newly predicted lncRNA was used as reference. Read counts computed by featureCounts were then subjected to differential expression analysis using edgeR [28]in R/Bioconductor environment.

### Reverse transcription and real-time quantitative PCR (qPCR)

cDNA synthesis was performed with one μg of total RNA in 20 μl reactions using the iScript Reverse Transcription Supermix (Bio-Rad Laboratories, California; cat. no. 1708840). qPCR was performed using iTaq™ Universal SYBR^®^ Green Supermix (Bio-Rad Laboratories, California; cat. no. 1725120), in a total of 20 μl reactions. All experiments were done in three biological and technical replicates. *RPS17* and *RPL32* were used as housekeeping genes, and all qPCR runs were carried out on the BioRad CFX96 qPCR platform. The 2-ΔΔCT method [29] was used to estimate the relative expression of each lncRNA, and the results were statistically analyzed using student’s t-tests. List of primers used in this study can be found in **S6 Table**.

### Functional annotation and enrichment

Functional annotation was performed using Pannzer [30] using amino acid sequences of lncRNA-target genes as input. Gene enrichment analysis was done using g;Profiler [31] and g:SCS threshold was used for multiple testing correction.

### CRISPR-Cas9 induced mutation of selected lncRNAs

sgRNA targeting exon 1 of lncRNA_27639.2 and XR_003899061.1 were designed using CHOPCHOP [32]. The sgRNAs were cloned into the *Drosophila* CRISPR-Cas9 vector pAc-dU6-sgRNA-Cas9 [33], which was a kind gift from Ji-Long Liu (Addgene plasmid 49330). This plasmid has been shown to work well in *Ae. aegypti* (Aag2) cell [34]. Detailed protocol for sgRNA cloning was previously described [33]. C6/36 cells (~3×10^6^) were transfected with 2.5 μg of gRNA-containing plasmid using Lipofectamine 2000 (Life Technology). Plasmid without gRNA (empty vector) was used as control. After 72 hours post-transfection, Cas9-expressing cells were selected using puromycin (5 μg/mL) for 5 days. Cells were then resuspended and ~5×10^5^ cells were added to each well of 12-well plate. Cells were allowed to settle for ~2 hour, before being infected with DENV-1 (Hawaiian strain) and ZIKV (H/PF/2013 strain) at MOI of 0.25. After 3 dpi, cells were frozen and thawed for twice, allowing the cells to break and release viral particles. The samples were then centrifuged, and the cell culture supernatant containing viral particles were used for TCID50 assay. Expression level of lncRNA after CRISPR-Cas9 induced mutation was evaluated by qPCR. RNA isolation and reverse transcription were carried out as previously described. Meanwhile, indel mutation within lncRNA loci was confirmed by sequencing of the PCR products spanning the cleavage sites. The PCR products were cloned into pGEM^®^-T Easy vector (Promega, cat. no: A1360), and ten colonies were selected for sequencing.

## Results

### Identification of novel lncRNAs in *Ae. albopictus*

To identify novel lncRNAs in *Ae. albopictus*, a total of 148 RNA-seq libraries were used as raw inputs. The list of all 148 RNA-seq datasets used in this study can be found in **S1 Data**. The RNA-seq libraries used were derived from many tissues, organs, developmental stages and cell lines; hence, allowing our lncRNA prediction to be more comprehensive. lncRNA prediction pipeline used here were modified and adapted from previous works [8], [9], [35]–[41]. Our lncRNA prediction pipeline (**Fig 1**) began with mapping the raw reads against the *Ae. albopictus* genome (AalbF2, assembly: GCA_006496715.1, NCBI) using Hisat2 [17], followed by transcriptome assembly by Stringtie [18]. The resulting transcriptome assemblies were merged into a single assembly, which was then compared to the reference annotation (Aalbo_primary.1_genomic.gtf, NCBI) using Gffcompare (https://github.com/gpertea/gffcompare). We selected transcripts with class code “i”, “u”, and “x” (35,863 transcripts), which indicate intronic, intergenic, and antisense to reference genes respectively [8], [40].

**Figure 1:**
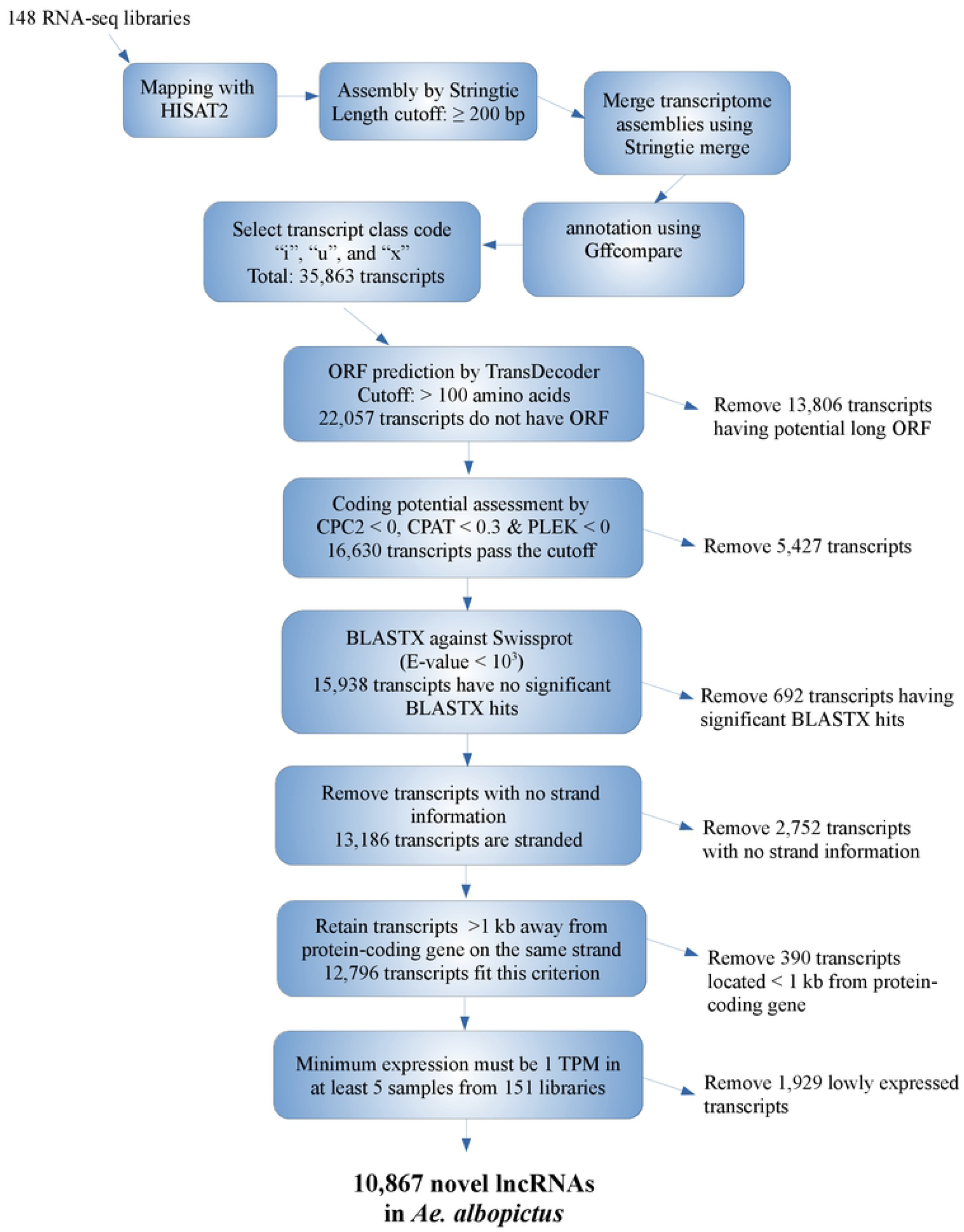
Overview of lncRNA identification pipeline

We then filtered the transcripts harboring open-reading frame (ORF) and coding potential. ORF prediction was performed using TransDecoder [19], while coding potential assessment was performed using three different algorithms, namely CPC2 [20], CPAT [22], and PLEK [21]. A total of 22,057 transcripts did not have ORF as determined from TransDecoder. Of these, 16,630 transcripts passed coding potential assessment by CPC2, CPAT, and PLEK. To avoid false positive prediction, we performed BLASTX on these 16,630 transcripts against Swissprot database. We removed transcripts having significant BLASTX hits (E-value < 10^3^). Next, we removed unstranded transcripts and those located less than 1 kb from protein-coding genes. Finally, we only retained transcripts that had minimum expression of 1 TPM in at least five out of the 148 RNA-seq libraries used in this study. This was done to remove lowly expressed transcripts which might be falsely identified as novel lncRNAs. In total, 10, 867 putative novel lncRNA transcripts, which derived from 9,298 genomic loci, were identified in *Ae. albopictus* genome. Of these 10,867 transcripts, 5,809 were located in the intergenic regions, 4,139 were intronic, while the remaining 919 were antisense to annotated genes. Full list of predicted novel lncRNAs and their genomic coordinates can be found in **S1 Table**. Current *Ae. albopictus* annotation catalogs 7,609 lncRNA transcripts, and we classified these transcripts as known lncRNAs. Therefore, in *Ae. albopictus* genome, the number of both novel and known lncRNAs were 18,476.

### *Ae. albopictus* lncRNAs shared similar features with other species

Next, we determined the similarities of *Ae. albopictus* lncRNAs to other species, such as shorter in length, low GC content, and low sequence conservation [8], [35], [36], [40]. *Ae. albopictus* lncRNAs were found to be significantly shorter in size compared to protein-coding transcripts (Kolmogorov-Smirnov test (KS-test) *p*-value < 2.2e-16, **Fig 2A**). The average length of novel and known lncRNAs in *Ae. albopictus* were 569.67 bp and 763.68 bp respectively, whereas protein-coding transcripts had a mean size of 2650.1 bp. Short in size is a typical feature of lncRNAs; hence, our findings are congruent with lncRNAs identified in other living organisms [8], [35], [42]. Besides, *Ae. albopictus* known and novel lncRNAs exhibited slightly lower GC content in comparison to protein-coding transcripts (**Fig 2B**). This lower GC or AT enriched characteristic is another notable feature of lncRNA transcripts [8], [9], [35], [36], [40]. Mean GC content of novel lncRNAs, known lncRNAs and protein-coding transcripts were 42%, 44%, and 48% respectively. We also analyzed GC content of other noncoding sequences in the genome such as 5’UTR and 3’UTR. We discovered that both of them had lower GC content in comparison to protein-coding transcripts. The average GC content of 5’UTR and 3’UTR were 44% and 36% respectively.

**Figure 2:**
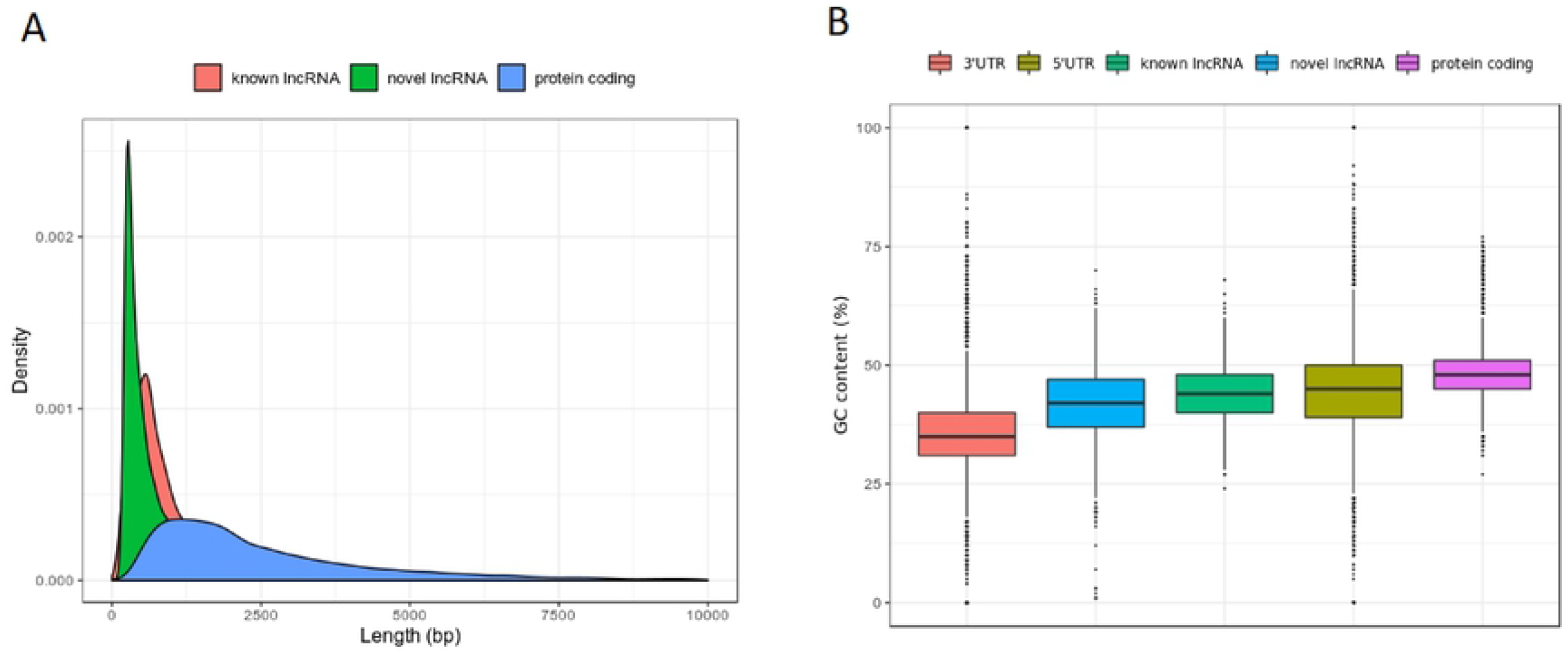
Length and GC content *Ae. albopictus* lncRNAs. A) Sequence length distribution of lncRNA and protein-coding transcript. B) GC content of lncRNA, protein-coding transcript, 3’UTR, and 5’UTR

As previous studies showed that lncRNAs demonstrate low sequence conservation even among closely related species [8], [9], [43], we investigated the validity of this in *Ae. albopictus*. To assess the level of sequence similarity, we performed BLASTN of *Ae. albopictus* lncRNAs against lncRNA transcripts found in other organisms including *Ae. aegypti, D. melanogaster, M. musculus, C. elegans*, and humans. lncRNA sequences of *Ae. aegypti* were taken from previous work [8], while lncRNA sequences of the remaining species were downloaded from NONCODE, a database for ncRNAs, especially lncRNAs [44]. BLAST algorithm bit score was used as an indicator for sequence similarity [8], [9]. In general, protein-coding transcripts were found to be more conserved than lncRNAs (**Fig 3A**).

**Figure 3:**
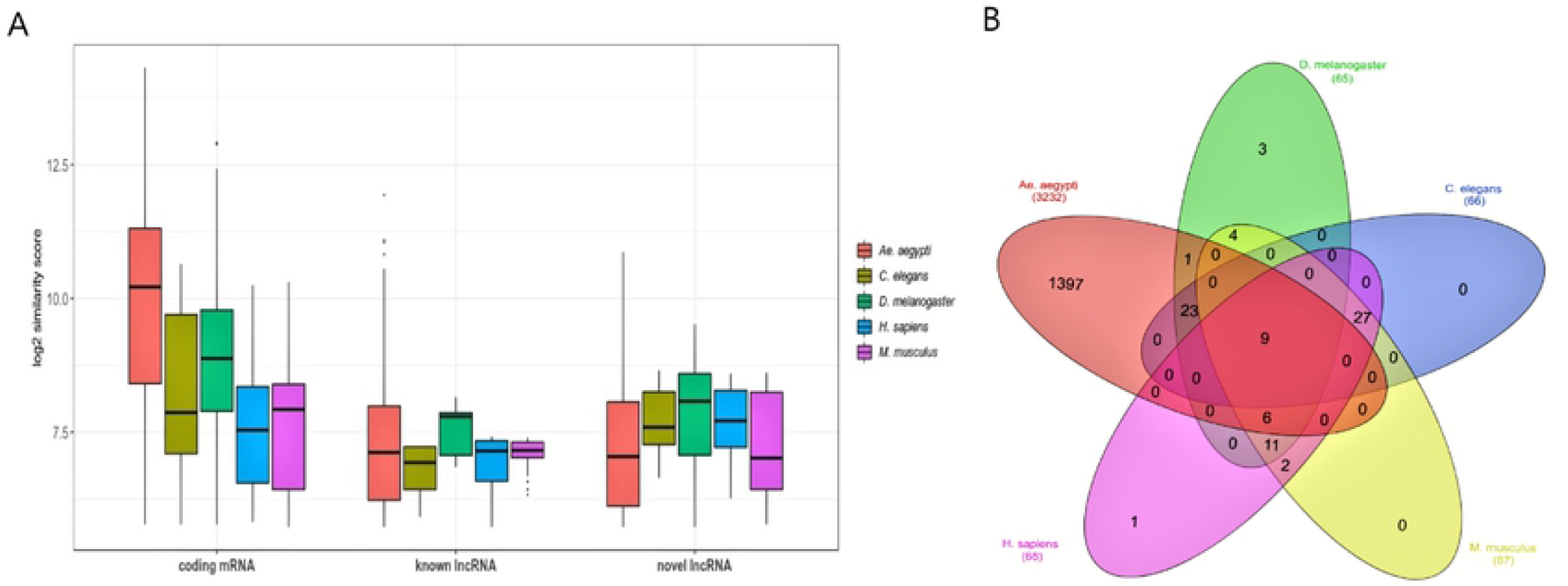
Conservation analysis of *Ae. albopictus* lncRNAs. A) Similarity bit score of of lncRNA and protein-coding mRNA. In general, protein-coding mRNA is more conserved than lncRNA. B) The Venn diagram shows the number of *Ae. albopictus* lncRNAs that display high conservation with other species according to BLASTN algorithm (E-value < 10-e5)

*Ae. albopictus* lncRNAs showed high degree of sequence similarity (E-value < 10^-5^) with *Ae. aegypti*, indicating that they are likely genus specific (**Fig 3B**). However, the fraction of *Ae. albopictus* lncRNAs that displayed high level of sequence similarity (E-value < 10^-5^) with *Ae. aegypti* were very low. For instance, only 819 (7.5%) novel lncRNAs and 617 (7.9%) known lncRNAs shared high sequence similarity with *Ae. aegypti*. On the other hand, 30,298 (75.6%) *Ae. albopictus* protein-coding transcripts demonstrated high degree of sequence similarity with *Ae. aegypti* protein-coding sequences.

### Overview of sequencing data generated from DENV and ZIKV-infected *Ae. albopictus* cells

Studies on *Ae. aegypti* transcriptomes provide evidence that lncRNAs could be involved in DENV and ZIKV-mosquito interaction [9], [45]. To examine the relevance of this to *Ae. albopictus*, we generated paired-end RNA-seq libraries from larval-derived *Ae. albopictus* cell line (C6/36) which had been infected with DENV and ZIKV. A total of 9 samples were sequenced using Illumina HiSeq 2000 platform, and they comprised of three replicates of mock, DENV, and ZIKV-infected C6/36 cells. In total, 426 million paired-end reads were generated in this study (**Table 1**). More than 70% of the reads (average of 35.5 million reads per sample) mapped to *Ae. albopictus* reference genome (**Table 1**), providing sufficient coverage for genome-wide transcriptomic analysis, specifically in detecting differentially expressed transcripts. Hierarchical clustering based on the expression values (CPM) of all the expressed genes showed that the replicates were grouped into their three distinct groups (**S1 Figure**). This unsupervised clustering also showed that the data formed two main clusters, i.e., uninfected, and virus-infected (DENV and ZIKV) samples.

### Differential expression of *Ae. albopictus* lncRNAs upon DENV and ZIKV infection

In this study, we considered genes with fold change (FC) of more than or equal to 2 in either direction, and false discovery rate (FDR) of less than 0.05 as differentially expressed. A total of 433 and 338 differentially expressed lncRNAs (FC ≥ |2|, FDR < 0.05) were identified respectively in DENV and ZIKV samples. In DENV-infected cells, 298 lncRNAs were found to be upregulated, while the remaining 136 transcripts were downregulated. Meanwhile, 216 and 122 lncRNAs were upregulated and downregulated respectively in C6/36 cells infected with ZIKV. Volcano plot analysis was used to depict the distribution of lncRNAs with their corresponding P-value and log FC (**Figure 4A**), and hierarchical clustering analysis was performed to cluster the differentially expressed lncRNAs among different replicates (**Figure 4B**). Similar lncRNAs were found to be differentially expressed −11 lncRNAs were upregulated and 12 lncRNAs were downregulated in both DENV and ZIKV infected cells (**Table 2**). The similar changes seen in these 23 lncRNAs post-infection suggest that they may be key players, regardless of the type of viral infection. On the other hand, the number of differentially expressed protein-coding mRNAs were higher, i.e., 1,718 and 1,098 mRNAs were differentially expressed (FC ≥ |2|, FDR < 0.05) in DENV and ZIKV-infected cells respectively. List of differentially expressed lncRNAs can be found in **S2 Table**.

**Figure 4:**
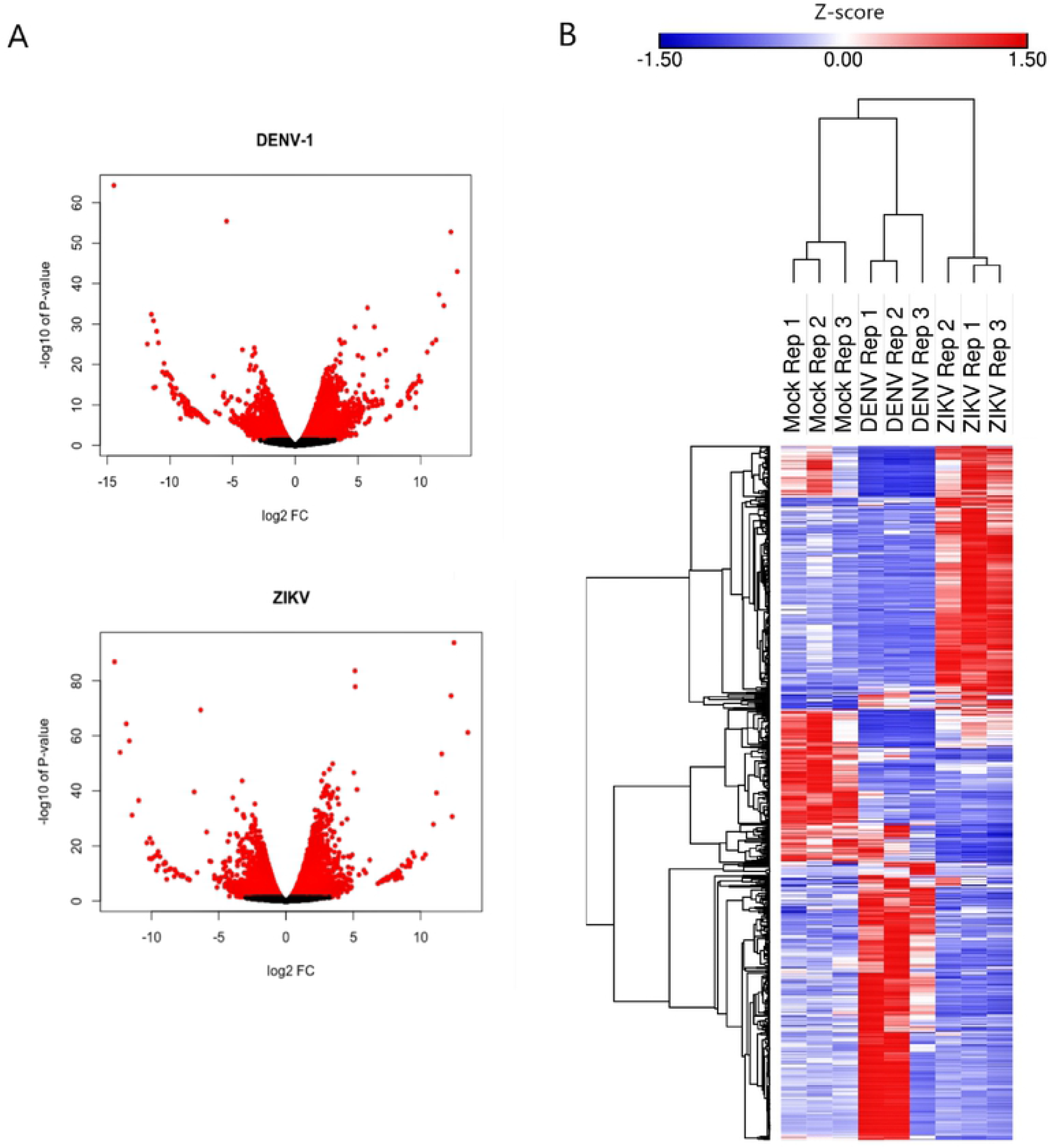
lncRNA expression profiles of DENV and ZIKV-infected *Ae. albopictus* cells. A) Volcano plot of differentially expressed *Ae. albopictus* lncRNAs in DENV-1 and ZIKV-infected C6/36 cells. Dots with red color represent lncRNAs with P-value < 0.05. B) Hierarchical clustering of differentially expressed *Ae. albopictus* lncRNA was done in Morpheus (https://software.broadinstitute.org/morpheus), using one minus Pearson correlation metric with average linkage method.

We randomly selected ten differentially expressed lncRNAs from RNA-seq analysis for qPCR. The ten chosen candidates were upregulated or downregulated in both DENV and ZIKV-infected samples. We validated the differentially expressed lncRNAs using qPCR from the same RNA samples used for Illumina sequencing. *RPS17* and *RPL32* were used as housekeeping genes [46], [47]. Our preliminary tests showed that the expression level of the two housekeeping genes were unchanged in DENV, and ZIKV-infected C6/36 cells; hence, qualifying them as suitable references in this study. Overall, the qPCR results of all the ten lncRNAs were consistent with RNA-seq data; hence, validating the bioinformatic approach used in this study (**Fig 5**)

**Figure 5:**
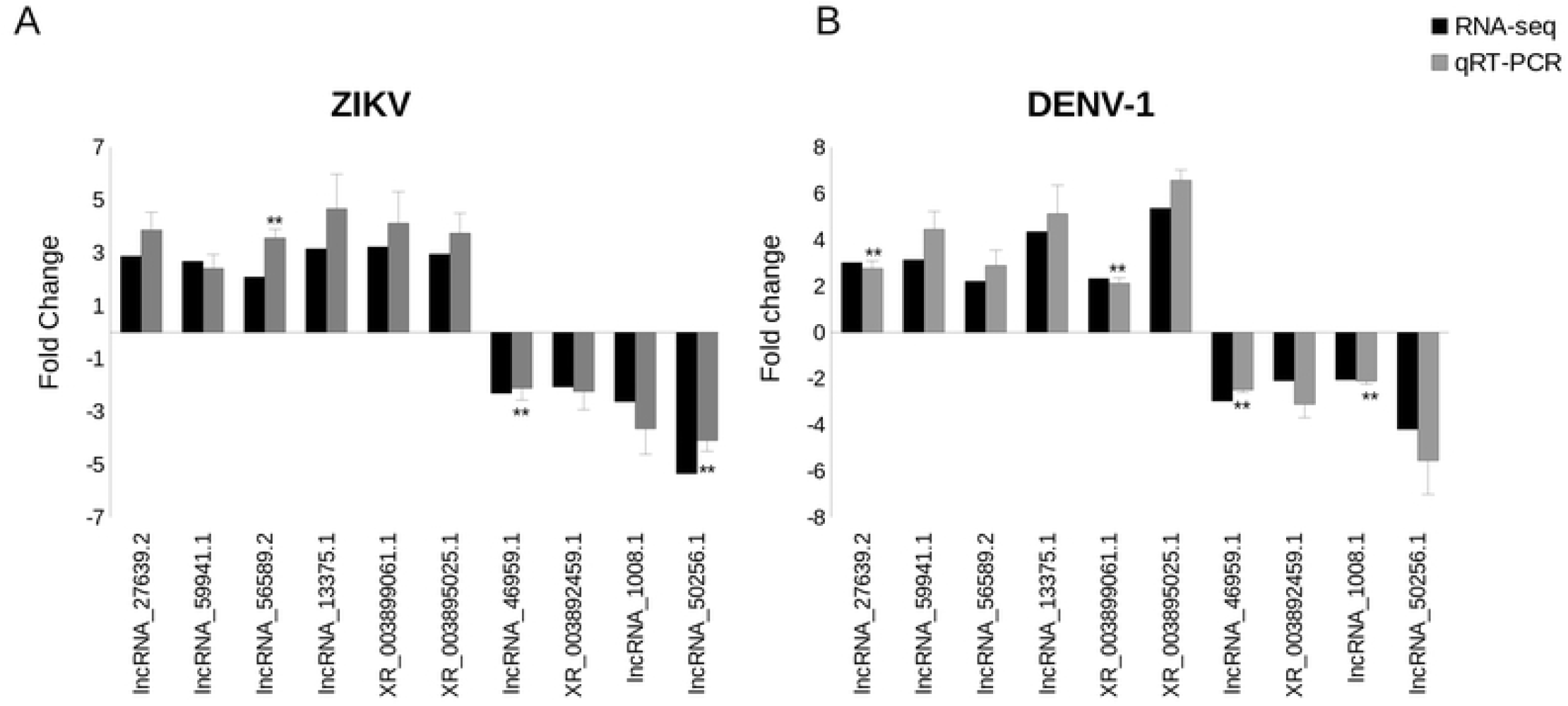
RT-qPCR validation of the differentially expressed lncRNAs. Expression levels were normalized to uninfected samples. Bars indicate mean +/- standard error of the mean (SEM) of three independent experiments. Student’s *t*-test was used to assess the statistical significance. All have p-value of < 0.05, and those of p-value < 0.01 are denoted with **. RNA-seq fold change was computed by edgeR.

### Functional analysis of the differentially expressed lnRNAs

To determine the putative regulatory roles of *Ae. albopictus* lncRNAs upon virus infections, we searched for protein-coding genes located within 100 kb (upstream and downstream) from differentially expressed lncRNAs as potential *cis*-regulated genes [48]–[50]and found a total of 198 and 153 coding genes which fulfilled that criteria in the DENV and ZIKV transcriptome respectively. As lncRNAs also have the ability to regulate genes in trans or distal regions [48], [51]–[53] we searched for protein-coding genes that were co-expressed with the differentially expressed lncRNAs based on Pearson correlation (coefficient > 0.95 or < −0.95, P-value < 0.05). In DENV transcriptome, 110 coding genes were co-expressed with the differentially expressed lncRNAs. Meanwhile, we detected 38 coding genes having high correlation in expression with differentially expressed lncRNAs in ZIKV transcriptome. We categorized the co-expressed genes as putative trans-regulated genes in *Ae. albopictus* cells upon DENV and ZIKV infection. Results of co-expression analysis between lncRNAs and protein-coding mRNAs can be found in **S3 Table**.

We selected the potential *cis* and *trans*-regulated genes for functional annotation and enrichment analyses. Previous reports demonstrated that upon virus infection in *Aedes* mosquitoes, there were changes in the expression level of host immune-related genes [13], [45], [54]–[56]. By conducting functional annotation using Pannzer [30] we discovered 13 lncRNAs that potentially regulated genes involved in immune-related functions (**S4 Table**). Based on gene ontology (GO), these genes were implicated in innate immune response (GO:0045087), defense response to Gram-positive (GO:0050830) and Gram-negative (GO:0050829), defense response to bacterium (GO:0042742), regulation of defense response to virus (GO:0050688), virus maturation (GO:0019075), and innate immune response (GO:0045087).

Previous studies reported that genes encoding clip-domain serine protease, and metalloproteinase were differentially expressed in *Aedes* mosquitoes upon virus infection. For example, clip-domain serine protease and metalloproteases were found to be significantly upregulated and downregulated respectively upon DENV infection in *Ae. albopictus* midguts [13]. Meanwhile, another study in *Ae. aegypti* reported that metalloproteinase and serine protease showed an increase in expression upon infection with ZIKV [45]. In this study, we discovered lncRNAs that potentially regulate genes that were also involved in serine-type endopeptidase activity (GO:004252, GO:004253, GO:004254) and metalloendopeptidase activity (GO:004222).

We then performed functional enrichment analysis of lncRNA-regulated genes using g;Profiler [31]. We discovered that 32 and 22 GO terms were significantly enriched (P-value < 0.05) in ZIKV and DENV-transcriptome respectively (Supplementary information 8, Supplementary Figure 2). A total of 8 GO terms were significant and shared by both transcriptomes, suggesting that lncRNAs may target genes that perform similar regulatory functions upon DENV and ZIKV infections (**S5 Table**). Among the significant GO terms were cellular processes, ion/protein binding, metabolic processes, and DNA binding.

### Mutation of selected lncRNAs and enhancement of DENV and ZIKV replication

To verify the role of *Ae. albopictus* lncRNAs in virus infection, we used CRISPR-Cas9 to induce mutation of selected lncRNAs in C6/36 cells following DENV and ZIKV infection. We discovered that CRISPR-Cas9 induced mutations in two lncRNAs loci resulted in the increase of DENV-1 and ZIKV infectious titer by at least two folds after three dpi (**Fig 6A**). One of the lncRNAs was novel (transcript ID: lncRNA_27639.2, loci ID: XLOC_029733) and another was known lncRNA (XR_003899061.1, ncbi gene ID: LOC115270134). Indels for both lncRNAs were confirmed by sequencing the PCR products spanning the cleavage sites, and the expression level lncRNAs after CRISPR-Cas9 induced mutation were validated by RT-qPCR (**S2 Figure**). Reduction in the level of expression indicated that mutations induced by CRISPR-Cas9 resulted in the silencing of the two aforementioned lncRNAs in C6/36 cells.

**Figure 6:**
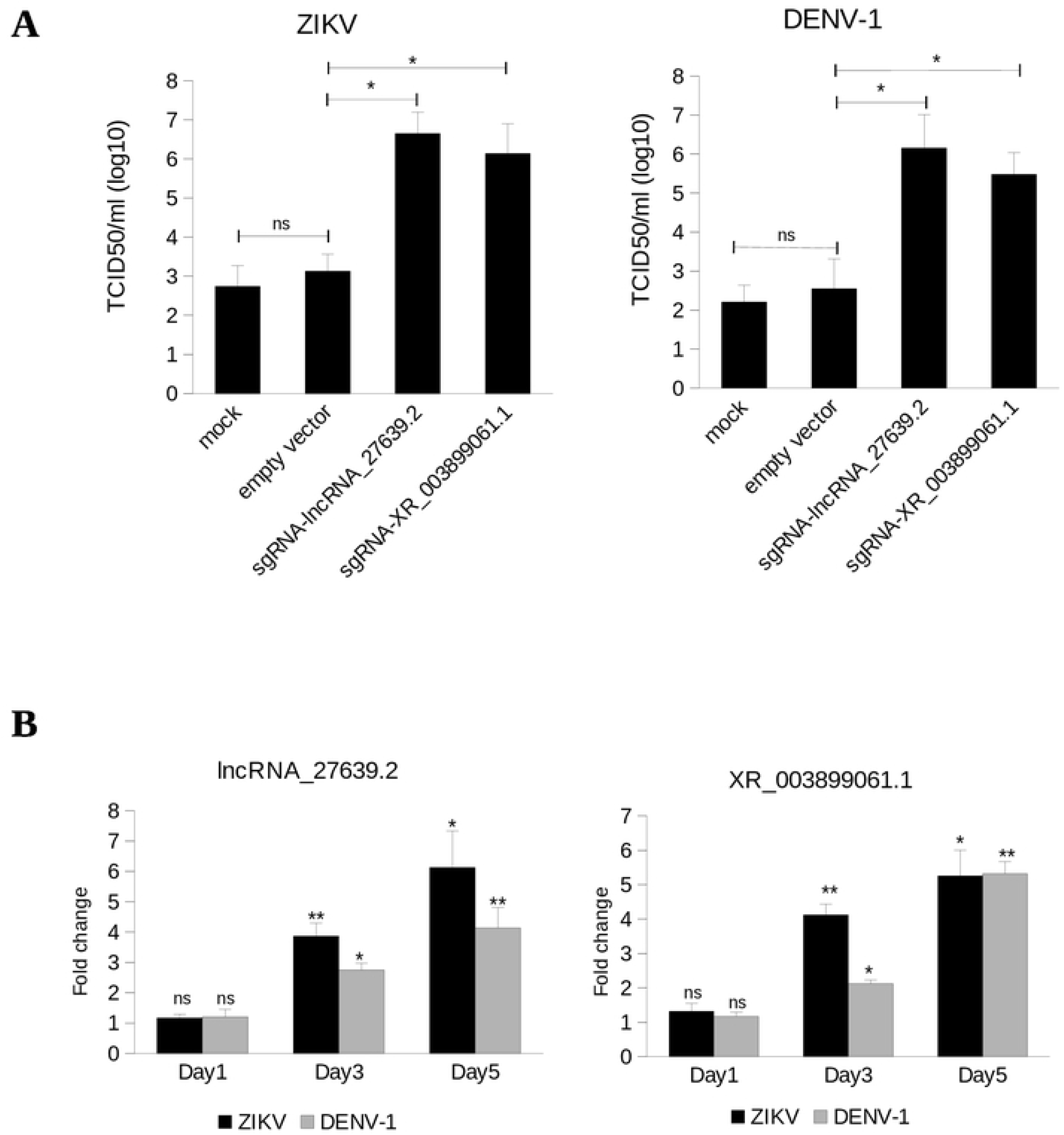
Possible involvement of lncRNAs in DENV and ZIKV replication in C6/36 cell. A) Titer of ZIKV and DENV-1 after 3 dpi as determined by TCID50 assay. In mock sample, C6/36 cells were transfected with transfection reagent only (Lipofectamine 2000). Empty vector refers to the *Drosophila* CRISPR-Cas9 vector (pAc-dU6-sgRNA-Cas9) without sgRNA targeting any gene. B) Expression level of lncRNA_27639.2 and XR_003899061.1 at 1, 3 and 5 dpi. At 1 dpi, the fold change between infected and uninfected samples were statistically insignificant. For A) and B), bars indicate mean +/- SEM of three independent experiments. Student’s *t*-test was used to assess the statistical significance. * p<0.05; **p<0.01.

A previous report showed that the expression of *Ae. aegypti* lincRNA_1317 increased substantially following the progression of infection [9]. When this lncRNA was silenced by RNAi, it led to an enhancement of DENV replication in *Ae. aegypti* cell, implying the involvement of certain lncRNAs in viral infection. We next analyzed our lncRNAs to idenfify if they also played a key role in anti-viral response. Based on RNA-seq and RT-qPCR analysis, both lncRNAs (lncRNA_27639.2 and XR_003899061.1) were upregulated following infection with DENV and ZIKV at 3 dpi (**Fig 5**). We investigated the level of expression at early (1 dpi) and late (5 dpi) stage of DENV and ZIKV infection. Interestingly, the expression of both lncRNAs displayed an increasing pattern along the progression of infection (**Fig 6B**), indicating possible involvement of lncRNA_27639.2 and XR_003899061.1 in defense against viruses in *Ae. albopictus*.

lncRNA_27639.2 might potentially regulate a gene that encodes for solute carrier family 35 member G1 (LOC109431891). This gene was located 1.9 kb downstream of lncRNA_27639.2 on the same strand; hence, making it a potential *cis*-regulated gene. GO annotation of solute carrier family 35 member G1 (LOC109431891) showed that it participates in calcium sensing and homeostasis, and it is a part of endoplasmic reticulum [57]. Meanwhile, XR_003899061.1, was located in *cis* to and 61 kb away from a gene encoding clathrin interactor 1 protein (LOC115269840), which is involved in clathrin assembly and trafficking [58]. Interestingly, calcium homeostasis, clathrin-mediated endocytosis, and vesicular trafficking are essential for viral replication in host cells [59]–[61]; thereby, further supporting the idea that lncRNA_27639.2 and XR_003899061.1 might be involved in antiviral response.

## Discussion

Even though protein-coding genes have always been the central focus in many host-virus interaction studies, recently, there are few investigations that provided evidence on the possible involvement of lncRNAs in *Aedes* mosquitoes, especially *Ae. aegypti* [9], [45]. Next-generation sequencing and bioinformatics analyses give scientists the opportunity to comprehensively identify lncRNAs in various species. Here, we provide a comprehensive annotation of novel lncRNAs in *Ae. albopictus* using the latest genome version (AalbF2). A previous study reported a total of 2,623 lncRNAs in *Ae. albopictus* [15]. Nonetheless, the identification was done using previous version of genome assembly (AaloF1). Due to the utilization of short-read sequencing and high levels of repetitive DNA sequences, AaloF1 assembly becomes highly fragmented and consists of more than 150, 000 scaffolds [1]. Therefore, in this study, we used the latest genome version, AalbF2 assembly [62], which had reduced scaffold number, and produced a substantially complete gene set with lower duplications as compared to the previous genomes of *Ae. albopictus* [1], [63], [64].

In this study, we provided a comprehensive list of lncRNAs in *Ae. albopictus* genome. We used a total of 148 RNA-seq libraries in our lncRNA prediction pipeline. The libraries were generated from various *Ae. albopictus* tissues, body parts, and developmental stages; hence, making a more comprehensive pipeline. Examination of genomic features showed that our newly predicted *Ae. albopictus* lncRNAs shared similar characteristics with lncRNAs from other species [8], [9], [35], [42], [43], [65], [66]. The characteristics include shorter in size, lower GC content, and lower sequence conservation as compared to protein-coding genes. These findings reflect the reliability of the lncRNA prediction pipeline used in this study. Shorter in length is a common feature of ncRNAs in many species, presumably due to the fact that, unlike protein-coding gene, lncRNAs are not required to have ORF, stop codon, 5’UTR and 3’UTR. Novel lncRNAs predicted in this study had lower GC content compared to protein-coding genes, which is in congruent with previous finding that concludes noncoding regions of the genome are usually composed of lower GC [67].

Unlike protein-coding genes, low sequence conservation of lncRNAs makes functional prediction challenging. However, there are research that suggests that lncRNAs could regulate the expression of their neighboring protein-coding genes [49]; thereby, enabling the prediction of the function of lncRNAs. Another widely used method that facilitates the prediction of lncRNA’s target is co-expression analysis by identifying protein-coding genes that exhibit similar expression patterns [48], [49] in trans or in distal regions. By identifying both cis and trans-regulated genes, we managed to functionally predict lncRNA functions following DENV and ZIKV infection in C6/36 cells.

Immune-related genes have always been implicated in host-virus interaction in *Aedes* mosquitoes [13], [55], [68]. Interestingly, our results on *Ae. albopictus* revealed a subset of upregulated lncRNAs that potentially regulated genes with GO terms related to immune response, or defense response to viruses and bacteria (GO:0050688, GO:0050830, GO:0050829, GO:0045087, GO:0042742, GO:0098586) (**S4 Table**). This implies an association between lncRNA overexpression and antiviral response. Functional enrichment analysis done in this study further supports the link between lncRNAs and virus infection, which is corroborated by previous studies in *Aedes* mosquitoes [45], [69], [70]. For example, GO terms associated with metabolic process, ion/protein binding, membrane-bounded organelle, and DNA binding were found to be significantly enriched in DENV and ZIKV infection, all of which have been identified in other transcriptomic studies in *Aedes* mosquitoes [45], [69], [70].

Reports on the association between silencing of host lncRNAs and replication of pathogens are still limited. One study reported that RNAi-mediated silencing of *Ae. aegypti* lincRNA_1317 led to enhancement of DENV replication [9]. A similar observation was made in HIV-1-infected human T cells, where the expression of lncRNA called NEAT1 was altered, and upon silencing of this lncRNA, the replication of virus was observed to be increased [71]. The depletion of two lncRNAs by CRISPR-Cas9 resulted in increased DENV and ZIKV infection in our studies, further validated the involvement of lncRNAs in virus infection. Therefore, this study provides strong evidence on the possible regulatory role of lncRNAs in virus infection of many host cells including humans.

Depletion of novel lncRNA, lncRNA_27639.2 (loci id: XLOC_029733), and known lncRNA, XR_003899061.1 (ncbi gene id: LOC115270134) in C6/36 cell led to enhancement of DENV-1 and ZIKV by at least two folds. Functional annotation revealed that lncRNA_27639.2 has a potential to regulate solute carrier family 35 member G1 (LOC109431891), whereas known lncRNA, XR_003899061.1, may regulate clathrin interactor 1 protein (LOC115269840). Solute carrier family 35 member G1 has been implicated in intracellular calcium sensing and homeostasis [57], a process that has been shown to be vital for virus replication [61]. For instance, infection of West Nile Virus (WNV) triggers an influx of calcium ion, and this process seems to be required for initial stage of infection and viral replication. This WNV-induced influx of calcium ion is probably necessary for rearrangement of the ER membrane to accommodate virus replication [72].

Even though there is no report that specifically links solute carrier family 35 member G1 to virus infection, its role in calcium homeostasis, which is important in virus replication, may implicate this gene in virus-host interaction. On the other hand, clathrin interactor 1 protein is essential for trafficking via clathrin-coated vesicles, and is localized at the trans-Golgi network [73]. Clathrin-mediated endocytosis is required for virus to enter cells including *Ae. albopictus* cells [59], [74], [75]. For example, treatment of C6/36 cells with chlorpromazine, an inhibitor of clathrin assembly, led to inhibition of DENV replication and entry [74]. Meanwhile, inhibition of clathrin-mediated endocytosis via RNAi-mediated silencing, or drug treatment (chlorpromazine and pitstop2) severely impaired ZIKV entry into human glioblastoma cell T98G cell [76].

Taken together, we provided a comprehensive genome-wide identification and characterization of *Ae. albopictus* lncRNAs and their involvement in DENV and ZIKV infection. Although our understanding on the functions of lncRNAs in *Aedes* mosquitoes is still limited, findings from this study may help future investigations. Further studies are required to functionally dissect the roles of lncRNA_27639.2 and XR_003899061.1 in DENV and ZIKV infection in not only *Ae. albopictus* cell, but in whole adult mosquitoes as well. We hypothesized that there are more lncRNAs related to antiviral response, so further characterization and functional studies should be done. We hope that this work will be an important resource for genetics and molecular studies of lncRNAs in *Ae. albopictus*.

## Acknowledgements

We would like to thank all our collaborators and colleagues for the discussion and the work conducted in this lab. This study was funded by the Universiti Sains Malaysia Research University Grant (1001/PBIOLOGI/811320 and 1001/PBIOLOGI/8011064) and ScienceFund Grant (305/PBIOLOGI/613238).

---

**Table 1: Summary of RNA-seq data**

**Table 2: Differentially expressed lncRNAs that are common in DENV-1 and ZIKV transcriptome** The value of logFC was determined by edgeR. All have FDR < 0.05

**S1 Data: List of 148 RNA-seq libraries used in this study**

**S1 Table: Genomic coordinates of *Ae. albopictus* novel lncRNAs**

**S2 Table: List of the differentially expressed lncRNAs**

**S3 Table: Results of co-expression analysis between lncRNAs and protein-coding genes**

**S4 Table: lncRNAs that potentially regulate immune-related genes**

**S5 Table: Significant GO terms found in DENV and ZIKV transcriptome**

**S6 Table: Primers used in this study**

**S1 Figure: Hierarchical clustering of RNA-seq samples** Hierarchical clustering was done using the Euclidean distance method associated with complete linkage

**S2 Figure: Validation of CRISPR-Cas9 induced mutation in lncRNA loci** A) As determined by RT-qPCR, CRISPR-Cas9 induced mutation resulted in lower expression level of lncRNA_27639.2 and XR_003899061.1 in C6/36 cells. Expression levels were normalized to empty vector sample. All changes in expression level shown in the figure are statistically significant with P-value < 0.05. Bars indicate mean +/- SEM of three independent experiments. Student’s *t*-test was used for all comparisons. B) Sequencing of indel mutations within lncRNA loci after 5 days of puromycin selection. PCR products flanking the cleavage site (black triangle) were cloned and sequenced from C6/36 cells transfected with vector containing lncRNA-specific sgRNAs. Majority of the clones showed deletions at the expected cleavage sites. Empty vector refers to the *Drosophila* CRISPR-Cas9 vector (pAc-dU6-sgRNA-Cas9) without gRNA targeting any gene. In mock sample, the C6/36 cells were transfected with transfection reagent only (Lipofectamine 2000). In both mock and empty vector samples, the clones showed wild type sequence. PAM is highlighted in blue, while target sequence in yellow.

